# Metabolic benefits of 17α-estradiol in liver are partially mediated by ERβ in male mice

**DOI:** 10.1101/2023.03.25.534216

**Authors:** Samim Ali Mondal, Shivani N. Mann, Carl van der Linden, Roshini Sathiaseelan, Maria Kamal, Snehasis Das, Matthew P. Bubak, Sreemathi Logan, Benjamin F. Miller, Michael B. Stout

**Affiliations:** Aging and Metabolism Research Program, Oklahoma Medical Research Foundation, Oklahoma City, OK, USA; Department of Neuroscience, University of Arizona, Tucson, AZ, USA; Department of Nutritional Sciences, University of Oklahoma Health Sciences Center, Oklahoma City, OK, USA; Department of Pathology, University of Oklahoma Health Sciences Center, Oklahoma City, OK, USA; Department of Physiology, University of Oklahoma Health Sciences Center, Oklahoma City, OK USA; Harold Hamm Diabetes Center, University of Oklahoma Health Sciences Center, Oklahoma City, USA; Department of Biochemistry & Molecular Biology, University of Oklahoma Health Sciences Center, Oklahoma City, OK USA; Oklahoma City Veterans Affairs Medical Center, Oklahoma City, OK, USA

## Abstract

Metabolic dysfunction underlies several chronic diseases. Dietary interventions can reverse metabolic declines and slow aging but remaining compliant is difficult. 17α-estradiol (17α-E2) treatment improves metabolic parameters and slows aging in male mice without inducing significant feminization. We recently reported that estrogen receptor α is required for the majority of 17α-E2-mediated benefits in male mice, but that 17α-E2 also attenuates fibrogenesis in liver, which is regulated by estrogen receptor β (ERβ)-expressing hepatic stellate cells (HSC). The current studies sought to determine if 17α-E2-mediated benefits on systemic and hepatic metabolism are ERβ-dependent. We found that 17α-E2 treatment reversed obesity and related systemic metabolic sequela in both male and female mice, but this was partially blocked in female, but not male, ERβKO mice. ERβ ablation in male mice attenuated 17α-E2-mediated benefits on hepatic stearoyl-coenyzme A desaturase 1 (SCD1) and transforming growth factor β1 (TGF-β1) production, which play critical roles in HSC activation and liver fibrosis. We also found that 17α-E2 treatment suppresses SCD1 production in cultured hepatocytes and hepatic stellate cells, indicating that 17α-E2 directly signals in both cell-types to suppress drivers of steatosis and fibrosis. We conclude that ERβ partially controls 17α-E2-mediated benefits on systemic metabolic regulation in female, but not male, mice, and that 17α-E2 likely signals through ERβ in HSCs to attenuate pro-fibrotic mechanisms.

## INTRODUCTION

Aging is the dominant risk factor for most chronic diseases, many of which are linked to tissue-specific metabolic perturbations^1,2^. Age-related declines in metabolic homeostasis are further exacerbated by obesity^3,4^, which has increased dramatically in older adults in recent decades^5,6^. Moreover, obesity is now recognized to exacerbate aging mechanisms and induce phenotypes more commonly observed with advancing age^7–13^. These observations have led to speculation that obesity may represent a mild progeria syndrome^2,14–17^. Although it is well-established that dietary interventions including chronic calorie restriction and various forms of fasting can reverse obesity- and age-related mechanisms that promote chronic diseases, many of these strategies are poorly tolerated due to adverse effects on mood, thermoregulation, and musculoskeletal mass^18,19^. These adverse health outcomes have fostered extensive investigation into pharmacological compounds that reverse metabolic dysfunction, attenuate aging mechanisms, and curtail chronic disease burden.

17α-estradiol (17α-E2) is one of the more recently studied compounds to demonstrate efficacy for beneficially modulating health outcomes in rodents. The NIA Interventions Testing Program (ITP) reported that 17α-E2 extends median lifespan in male, but not female, mice when treatment is initiated in mid-life^20,21^ and late-life^22^. The magnitude of lifespan extension with 17α-E2 treatment is similar to that of calorie restriction^23^ and rapamycin administration^24^ in male mice. We have shown that 17α-E2 administration reduces calorie intake and regional adiposity in combination with significant improvements in several metabolic measures including glucose tolerance, insulin sensitivity, and ectopic lipid deposition in obese and aged male mice^25–29^. Other groups have also reported that 17α-E2 treatment elicits similar benefits on glucose tolerance, mTORC2 signaling, hepatic urea cycling, markers of neuroinflammation, and sarcopenia^30–34^. Importantly, male-specific benefits occur without significant feminization of the sex hormone profiles^25^ or reproductive function^35^. Female mice with intact ovarian function are generally unresponsive to 17α-E2 treatment^30–34,36,37^, although female mice with established metabolic dysfunction are yet to be tested. Ovariectomy renders female mice more responsive to the metabolic benefits of 17α-E2 treatment^38^, but chronic administration in ovariectomized females does not appear to curtail pro-aging mechanisms similarly to what is observed in male mice^30–34^. In contrast to female mice, intact female rats appear more responsive to 17α-E2 treatment^39^, although the mechanisms underlying this phenomenon remain unexplored.

Until recently the receptor(s) that mediate the actions of 17α-E2 were believed to be uncharacterized^40–43^ due to the relatively low binding affinity for classical estrogen receptors (ERα & ERβ) when compared to 17β-estradiol (17β-E2)^44,45^. However, we recently demonstrated that 17α-E2 and 17β-E2 elicit nearly identical genomic actions through ERα in an *in vitro* system and that the global ablation of ERα attenuates nearly all the metabolic benefits of 17α-E2 in male mice^27^. These observations indicate that 17α-E2 signals through ERα to elicit health benefits. This report also provided strong evidence that liver is one of the primary organs where 17α-E2 signals to modulate systemic metabolism^27^. Our subsequent study using a liver injury model revealed that 17α-E2 also attenuates fibrogenesis in liver^46^, which is dominantly mediated by hepatic stellate cells (HSCs). Interestingly, HSCs almost exclusively express ERβ^47,48^, which led us to hypothesize that 17α-E2 may also be eliciting partial metabolic benefits through ERβ, particularly within liver.

In the current study, we sought to determine if the global ablation of ERβ would curtail 17α-E2-mediated benefits on systemic metabolism in diet-induced obese mice that had been subjected to chronic high-fat diet (HFD) feeding prior to study initiation. We chose to challenge our mice for an extended period of time (9 months) prior to study initiation because it would enable us to also determine if female mice with established metabolic dysfunction would become responsive to 17α-E2 treatment. We found that ERβ is not required for 17α-E2 to elicit systemic metabolic benefits in male mice. We also found that chronically challenged female mice do indeed benefit from 17α-E2 treatment and that ERβ partially mediates these effects. Similar to our previous reports, we also found that 17α-E2 significantly improves readouts related to liver steatosis and fibrosis to varying degrees in both sexes. However, ERβ ablation in male mice partially attenuated 17α-E2-mediated benefits on hepatic stearoyl-coenyzme A desaturase 1 (SCD1) and transforming growth factor β1 (TGF-β1) production; both of which play a role in HSC activation and liver fibrosis^49–51^. Therefore, we subsequently evaluated the effects of 17α-E2 treatment *in vitro* on hepatocytes (HepG2) and hepatic stellate cells (LX-2) and found that 17α-E2 directly mitigates SCD1 production in both cell-types, thereby indicating that 17α-E2 directly signals in both cell-types to suppress drivers of steatosis and fibrosis. We conclude that ERβ does partially control 17α-E2-mediated benefits on the regulation of body mass and adiposity in female, but not male mice, and that 17α-E2 likely signals through ERβ in HSCs to attenuate pro-fibrotic mechanisms.

## RESULTS

### ERβ ablation affects metabolic responsiveness to 17α-E2 treatment in a sex-specific fashion

To induce obesity and metabolic perturbations in our mice, we administered HFD for 9 months prior to initiating 17α-E2 treatment. Immediately after 17α-E2 treatment began, male WT mice displayed significant reductions in mass (Fig. 1A,B) and adiposity (Fig. 1C,D), which is aligned with our previous reports^25–29^. Interestingly, male ERβKO mice responded almost identically to male WT mice with regard to reductions in mass and adiposity (Fig. 1A-D), indicating that ERβ is not involved in 17α-E2-mediated reductions in body mass and adiposity. Lean mass was unchanged by 17α-E2 in male mice of either genotype (Suppl. Fig. 1A). Unexpectedly, 17α-E2 treatment did not reduce calorie consumption in male WT or ERβKO mice (two-way ANOVA) in this study (Fig. 2A), although intake was suppressed with 17α-E2 treatment in both genotypes during the first two weeks of the intervention (p-values ranged from 0.026 – 0.078 when assessed by t-test within genotype across treatment groups). Both male WT and ERβKO mice receiving 17α-E2 displayed modest improvements in glucose tolerance (Fig. 3A,B), although this did not reach statistical significance. Conversely, insulin tolerance was significantly improved by 17α-E2 treatment in both male WT and ERβKO mice (Fig. 3C,D). 17α-E2 treatment also dramatically suppressed hyperinsulinemia in both male WT and ERβKO mice, providing additional evidence of improvements in metabolic health (Suppl. Fig. 2A).

**Figure 1.**
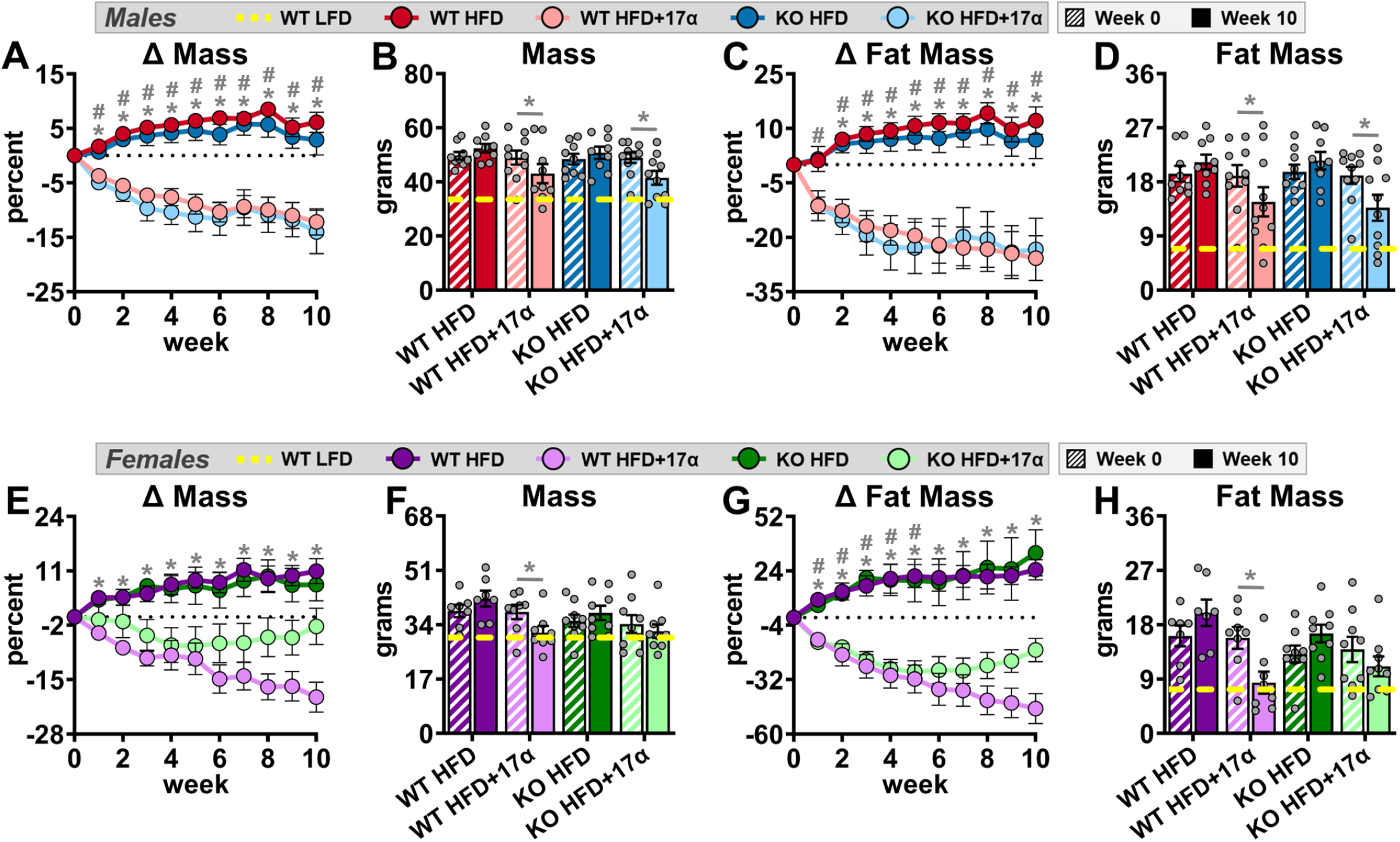
ERβ partially mediates 17α-E2 effects on mass and adiposity in obese female, but not male, mice. (A) Percent change in body mass over time [n=9-10/group/timepoint], (B) Body mass at baseline (week 0; striped) and week 10 (solid) [n=9-10/group/timepoint], (C) Percent change in fat mass over time [n=9-10/group/timepoint], and (D) Fat mass at baseline (week 0; striped) and week 10 (solid) [n=9-10/group/timepoint] in male WT and ERβKO mice. (E) Percent change in body mass over time [n=7-9/group/timepoint], (F) Body mass at baseline (week 0; striped) and week 10 (solid) [n=7-9/group/timepoint], (G) Percent change in fat mass over time [n=7-9/group/timepoint], and (H) Fat mass at baseline (week 0; striped) and week 10 (solid) [n=7-9/group/timepoint] in female WT and ERβKO mice. Age-matched, WT, LFD-fed mice were also evaluated as a normal-weight reference group and their corresponding means for both sexes are depicted as dashed yellow lines [n=9/group/timepoint]. All data are presented as mean ± SEM and were analyzed within sex by two-way repeated measures ANOVA with Tukey post-hoc comparisons. For panels A, C, E, & G, * represents differences between WT HFD and WT HFD+17α-E2, while # represents differences between ERβKO HFD and ERβKO HFD+17α-E2. For panels B, D, F, & H, * represents differences within treatment group over time. *#p < 0.05.

**Figure 2.**
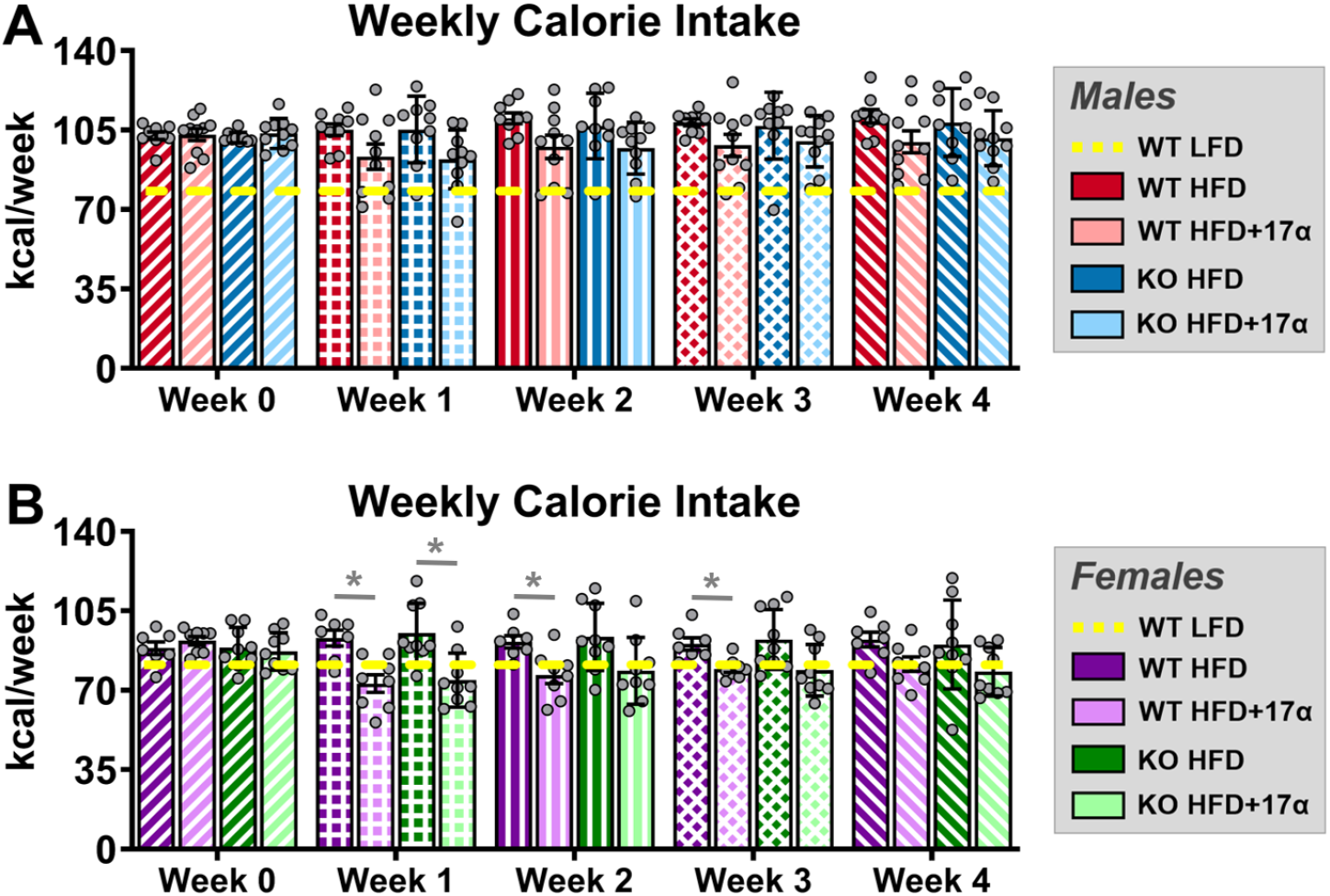
ERβ partially mediates 17α-E2 effects on calorie consumption in obese female, but not male, mice. (A) Average weekly calorie consumption during baseline (week 0) and throughout the first 4 weeks of treatment in male WT and ERβKO mice [n=9-10/group/timepoint]. (B) Average weekly calorie consumption during baseline (week 0) and throughout the first 4 weeks of treatment in female WT and ERβKO mice [n=7-9/group/timepoint]. Age-matched, WT, LFD-fed mice were also evaluated as a normal-weight reference group and their corresponding means for both sexes are depicted as dashed yellow lines [n=9/group/timepoint]. All data are presented as mean ± SEM and were analyzed within sex by two-way repeated measures ANOVA with Tukey post-hoc comparisons. * represents differences within genotypes across treatment groups at each timepoint. *p < 0.05.

**Figure 3.**
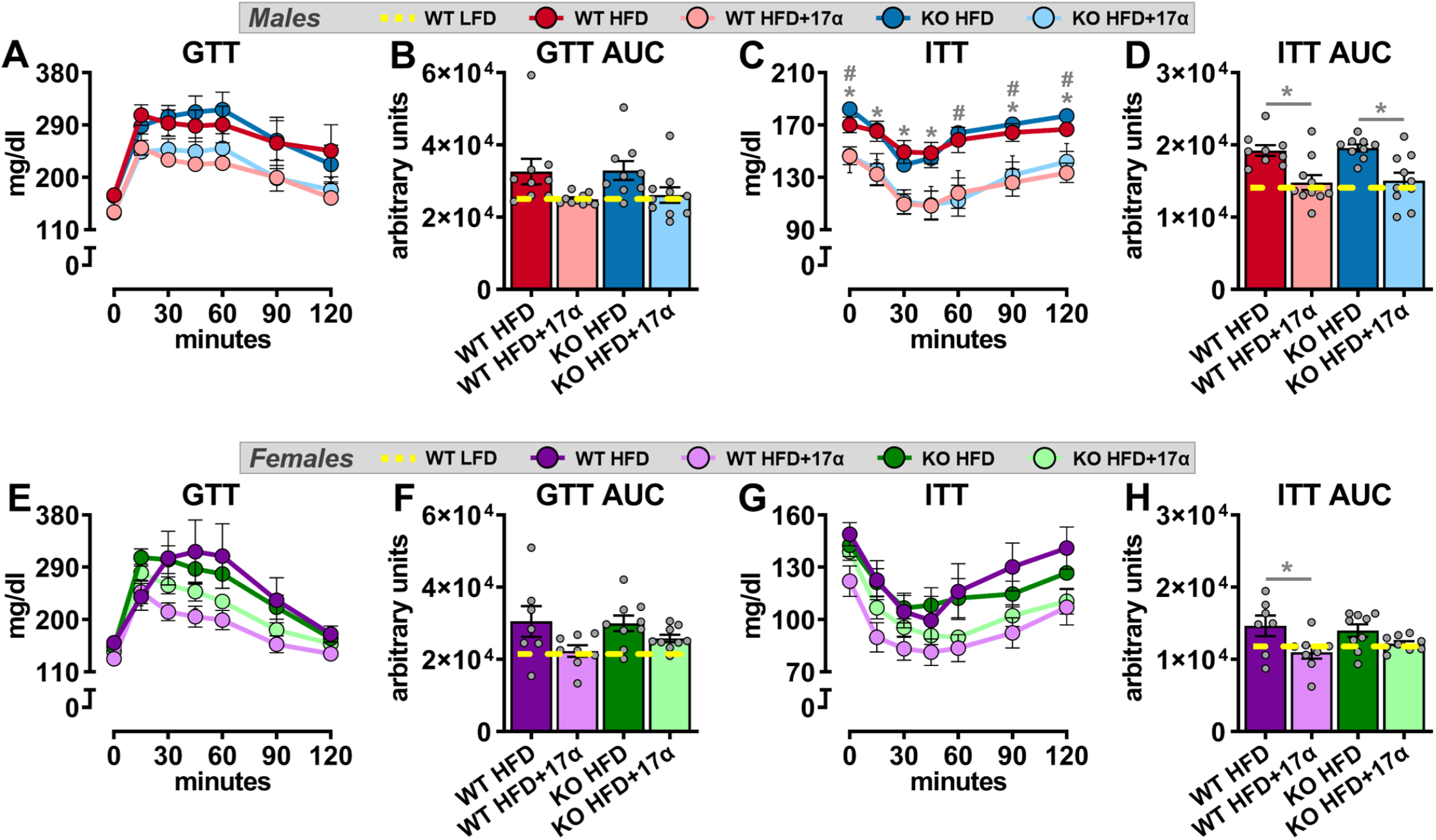
ERβ partially mediates improvements in insulin responsiveness with 17α-E2 treatment in obese female, but not male, mice. (A) GTT [n=9-10/group] and (B) GTT AUC [n=9-10/group] during week 9 in male WT and ERβKO mice. (C) ITT [n=9-10/group] and (D) ITT AUC [n=9-10/group] during week 10 in male WT and ERβKO mice. (E) GTT [n=7-9/group] and (F) GTT AUC [n=7-9/group] during week 9 in female WT and ERβKO mice. (G) ITT [n=7-9/group] and (H) ITT AUC [n=7-9/group] during week 10 in female WT and ERβKO mice. Age-matched, WT, LFD-fed mice were also evaluated as a normal-weight reference group and their corresponding means for both sexes are depicted as dashed yellow lines [n=9/group]. All data are presented as mean ± SEM and were analyzed within sex by two-way repeated measures ANOVA (A, C, E, G) or two-way ANOVA (B, D, F, H) with Tukey post-hoc comparisons. For panel C, * represents differences between WT HFD and WT HFD+17α-E2, while # represents differences between ERβKO HFD and ERβKO HFD+17α-E2. For panels D & H, * represents differences within genotypes across treatment groups. *p < 0.05.

In contrast to previous reports^30–34,36,37^, 17α-E2 treatment robustly reduced mass (Fig. 1E,F) and adiposity (Fig. 1G,H) in female WT mice in our study. Conversely, female ERβKO mice did not significantly reduce body mass in response to 17α-E2 treatment (Fig. 1E,F) and only initially reduced adiposity, but this effect waned over time (Fig. 1G,H). These were unexpected findings that indicate ERβ at least partially mediates the loss of body mass and adiposity attributed to 17α-E2 in obese female mice. As seen in males, lean mass was also unaffected by 17α-E2 in female mice of either genotype (Suppl. Fig. 1B). Similar to changes in adiposity, 17α-E2 treatment significantly reduced calorie consumption in female WT mice over the first three weeks of treatment, but this reduction only occurred during the first week of treatment in female ERβKO mice (Fig. 2B). Similar to males, both female WT and ERβKO mice receiving 17α-E2 displayed only mild, nonsignificant improvements in glucose tolerance (Fig. 3E,F). Interestingly, 17α-E2 treatment only improved insulin tolerance in female WT, and not ERβKO, mice (Fig. 3G,H). Genotype-specific responsiveness to 17α-E2 treatment in female mice was also observed in fasting insulin levels (Suppl. Fig. 2B), which only improved in the WT mice receiving 17α-E2. These observations suggest that ERβ is required for 17α-E2 to improve insulin sensitivity in obese female mice.

### 17α-E2 reverses obesity-related hepatic steatosis and other markers of liver disease independently of ERβ

We previously demonstrated that 17α-E2 suppresses hepatic lipid deposition in male mice through what appear to be a variety of mechanisms^25,27,46^. In the current study we sought to determine if these benefits require ERβ. We found that 17α-E2 significantly reduced liver mass and steatosis in both male WT and ERβKO mice (Fig. 4A,B). Pathological assessment confirmed the reversal of hepatic steatosis with 17α-E2 treatment in both genotypes (Fig. 4C,D). Interestingly, the benefits on steatosis were not associated with significant changes in transcriptional markers of hepatic lipogenesis (*Srebp1, Fasn, Acc1*; Suppl. Fig. 3A-C), although we are not the first to report this observation^52^. Pathological assessment also confirmed modest reductions in liver inflammation with 17α-E2 treatment in both male genotypes (Fig. 4C), although this did not rise to the level of significance. Subsequent evaluation of liver macrophage content and polarity supported the pathological data by revealing that 17α-E2 treatment did not significantly alter total macrophage (F4/80), M1 pro-inflammatory macrophage (CD11c), or M2 anti-inflammatory macrophage (CD206) content in either genotype (Suppl. Fig. 4A-D). We surmise that the lack of changes in liver macrophage outcomes in males receiving 17α-E2 stems from the fact that liver injury is still fairly minimal in this study. However, 17α-E2 treatment did significantly suppress liver *TNFα* transcription in male WT and ERβKO mice (Suppl. Fig. 5A), which is an important driver of liver disease^53^.

**Figure 4.**
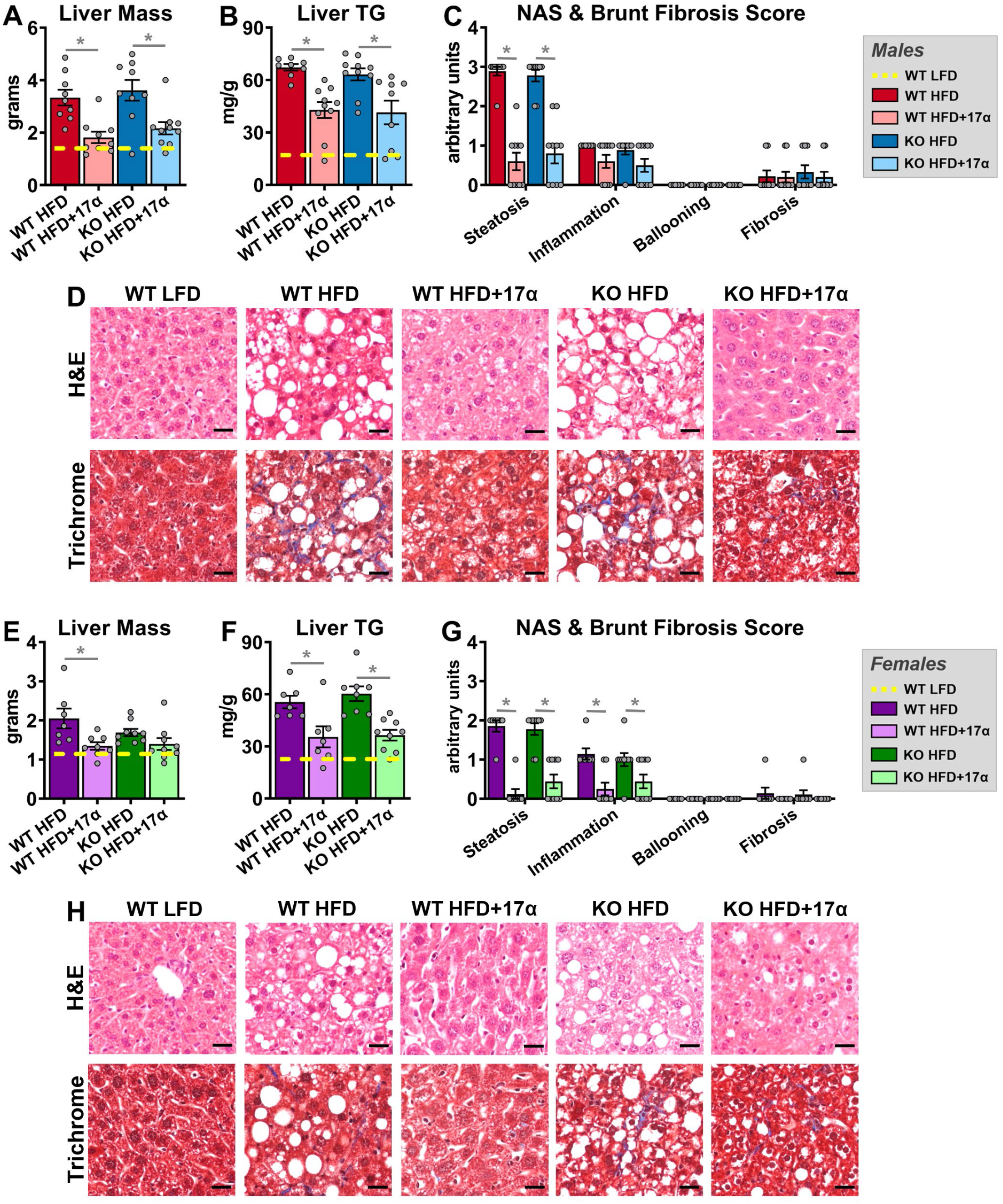
17α-E2 reverses obesity-related hepatic steatosis in both sexes in an ERβ-independent manner. (A) Liver mass [n=9-10/group], (B) Liver TG [n=9-10/group], (C) Liver pathological NAS and Brunt Fibrosis Scores [n=9-10/group], and (D) Representative images of gross morphology, H&E stained (magnification = 20X; scale bar = 50 μm) and Masson’s trichrome stained (magnification = 20X; scale bar = 50 μm), liver from male WT and ERβKO mice. (E) Liver mass [n=7-9/group], (F) Liver TG [n=7-9/group], (G) Liver pathological NAS and Brunt Fibrosis Scores [n=7-9/group], and (D) Representative images of gross morphology, H&E stained (magnification = 20X; scale bar = 50 μm) and Masson’s trichrome stained (magnification = 20X; scale bar = 50 μm), liver from female WT and ERβKO mice. Age-matched, WT, LFD-fed mice were also evaluated as a normal-weight reference group and their corresponding means for both sexes are depicted as dashed yellow lines [n=9/group/timepoint], except for liver NAS and Brunt Fibrosis Scores because all scores were zero. All data are presented as mean ± SEM and were analyzed within sex by two-way ANOVA with Tukey post-hoc comparisons. * represents differences within genotypes across treatment groups. *p < 0.05.

In females, 17α-E2 treatment reduced liver mass only in WT mice (Fig. 4E), but reduced liver steatosis in both genotypes (Fig. 4F); the latter of which was confirmed by pathological assessment (Fig. 4G,H). Similar to the findings in males, these benefits were not associated with significant changes in transcriptional markers of hepatic lipogenesis (Suppl. Fig. 3D-F). In contrast to males, pathological assessment revealed that 17α-E2 treatment significantly reduced liver inflammation in both female WT and ERβKO mice (Fig. 4G). Although 17α-E2 treatment failed to reduce liver total macrophage (F4/80) content in either genotype (Suppl. Fig. 4E,F), it did significantly suppress M1 pro-inflammatory macrophage (CD11c) content in female WT mice (Suppl. Fig. 4E, G), which is aligned with the previously addressed histopathological data. M2 anti-inflammatory macrophage (CD206) content was also unchanged by 17α-E2 treatment in females of either genotype (Suppl. Fig 4E,H). As expected, hepatic *TNFα* transcripts were suppressed by 17α-E2 treatment in obese female mice, but this only rose to the level of statistical significance in the ERβKO mice (Suppl. Fig. 5B).

### 17α-E2 treatment suppresses mechanistic drivers of liver disease in an ERβ-dependent manner in male mice

In alignment with declines in hepatic steatosis, we also found that 17α-E2 treatment robustly downregulated hepatic SCD1 protein in male WT, but not ERβKO, mice (Fig. 5A,B), which is congruent with our prior report^46^. Although the magnitude of liver fibrosis in this study was mild (Fig. 4C,D), which is common in mice when HFD is the sole challenge^54^, we still evaluated the underlying drivers of hepatic fibrogenesis due to our prior report demonstrating that 17α-E2 reduced liver fibrosis in a model of liver injury^46^. We found that 17α-E2 treatment significantly suppressed hepatic TGF-β1 production in WT mice, but this was prevented by ERβ ablation (Fig. 4C). TGF-β1 is the dominant mechanistic driver of liver fibrosis^49,50^ and is known to be produced in both HSCs and macrophages during liver injury^49,55,56^. The absence of significant 17α-E2-mediated effects on hepatic SCD1 and TGF-β1 production in male ERβKO mice suggests that 17α-E2 likely elicits benefits in cell-types that predominantly express ERβ, which include HSCs^47,48^. The effects of 17α-E2 treatment on hepatic SCD1 protein in female mice mirrored our findings in male mice, with only WTs displaying improvements (Fig. 5D,E). It should be noted that 17α-E2 did moderately suppress hepatic SCD1 in female ERβKO mice, but this did not reach statistical significance. In contrast to males, 17α-E2 treatment failed to modulate hepatic TGF-β1 production in female mice of either genotype (Fig. 5F). These observations are highly suggestive that 17α-E2 suppresses SCD1 in hepatocytes as a means of curtailing hepatic lipid accumulation in female mice, but that 17α-E2 is either incapable of modulating pro-fibrotic mechanisms in female liver, or the female mice in our study were too metabolically healthy to develop sufficient liver pathology for 17α-E2 to elicit benefits.

**Figure 5.**
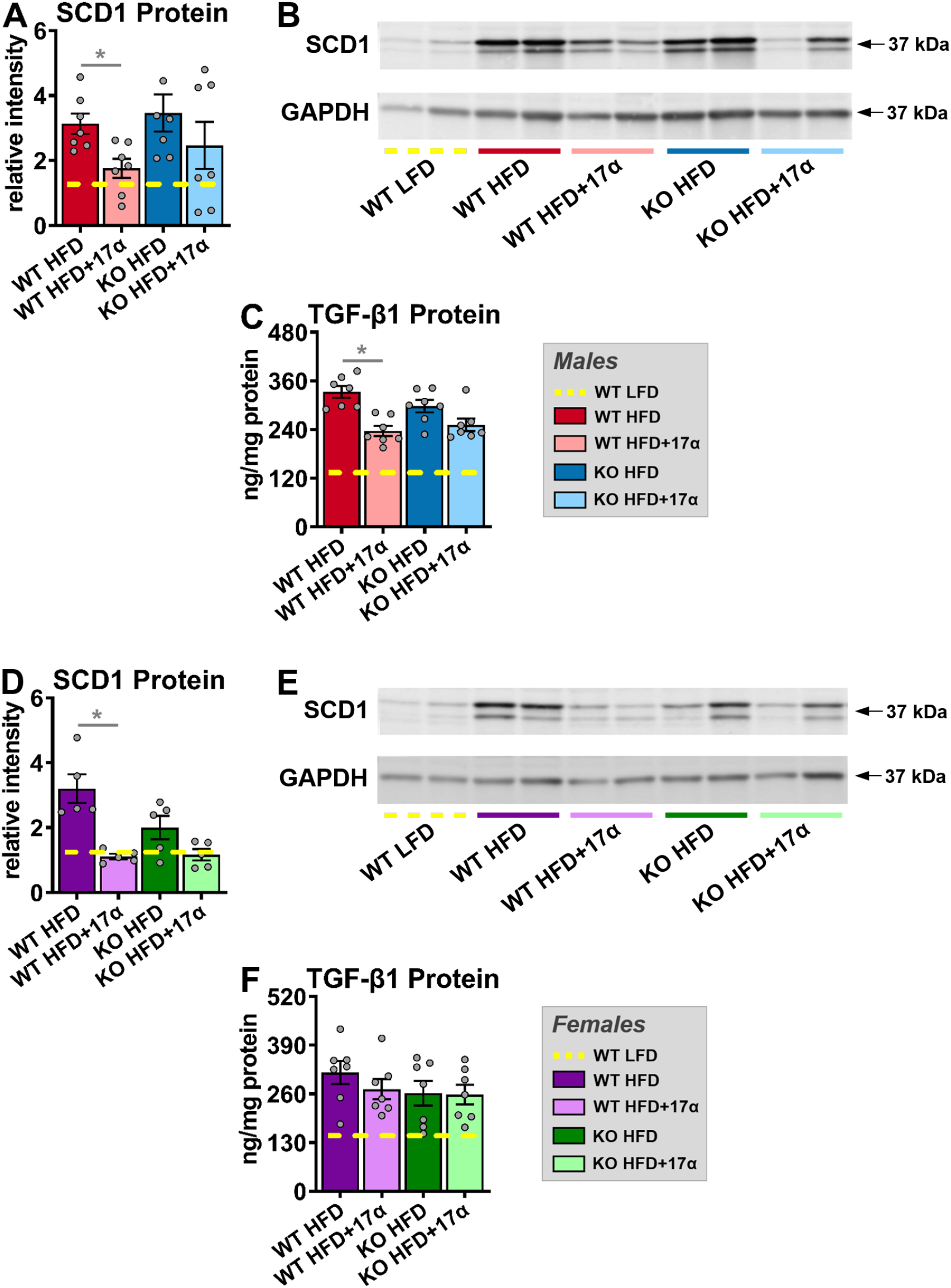
ERβ partially regulates 17α-E2-mediated effects on markers associated with hepatic steatosis and fibrosis in a sex-specific manner. (A) SCD1 protein [n=7/group], (B) Representative immunoblots of SCD1 and GAPDH, and (C) TGF-β1 protein [n=7/group] in liver from male WT and ERβKO mice. (D) SCD1 protein [n=5/group], (E) Representative immunoblots of SCD1 and GAPDH, and (F) TGF-β1 protein [n=7/group] in liver from female WT and ERβKO mice. Age-matched, WT, LFD-fed mice were also evaluated as a normal-weight reference group and their corresponding means for both sexes are depicted as dashed yellow lines [n=5-7/group]. All data are presented as mean ± SEM and were analyzed within sex by two-way ANOVA with Tukey post-hoc comparisons. * represents differences within genotypes across treatment groups. *p < 0.05.

### 17α-E2 attenuates SCD1 production in both hepatocytes and HSCs *in vitro*

In an effort to determine if 17α-E2 directly modulates SCD1 production in hepatocytes and HSCs, we performed *in vitro* studies using HepG2 and LX-2 cells that had been challenged with PA and TGF-β1, respectively. We initially performed time course evaluations to ensure that adequate survival of HepG2 cells were maintained following PA treatment, and proliferation (*i.e*. activation) of LX-2 cells occurred following TGF-β1 treatment. We found that 6 and 12 hours of PA treatment were ideal for HepG2 viability (> 75%), whereas 24 hours of treatment significantly reduced HepG2 viability (< 60%) (Fig. 6A). These observations were supported by increased *Bax* transcription, a marker of apoptosis^57^, only following 24 hours of PA treatment (Fig. 6B). In LX-2 cells, we found that 12 and 24 hours of TGF-β1 treatment induced significant proliferation (≥ 40%), which is common when HSCs are activated^49,50^, whereas 6 hours of treatment failed to induce proliferation (Fig. 6C). Collagen transcription was also increased following 12 and 24, but not 6, hours of TGF-β1 exposure (Fig. 6D). The findings outlined above provided the justification to perform our subsequent HepG2 experiments following 6 and 12 hours of PA exposure, and our LX-2 experiments following 12 and 24 hours of TGF-β1 exposure. We then evaluated *Scd1* mRNA induction in HepG2 cells treated with PA and found that 12 hours of treatment significantly increased *Scd1* transcription, and that this was completely attenuated by 1 nM of 17α-E2 co-treatment (Fig. 6E). To determine if this finding translated to the protein level, we then evaluated SCD1 protein following 12 hours of PA treatment and found it was indeed significantly upregulated, but that all dosing regimens (100 nM, 10 nM, 1 nM) of 17α-E2 co-treatment curtailed this induction (Fig. 6F,G). These findings clearly indicate that 17α-E2 can suppress SCD1 expression in stressed hepatocytes, which was anticipated due to our previous reports^25,27,46^. We then evaluated *Scd1* mRNA induction in LX-2 cells treated with TGF-β1 and found that both 12 and 24 hours of treatment significantly increased *Scd1* transcription, but that 12 hours displayed a more robust induction of *Scd1* mRNA and that this was almost completely attenuated by 10 nM and 1 nM of 17α-E2 co-treatment (Fig. 6H). We then evaluated SCD1 protein following 12 hours of TGF-β1 treatment and found it was mildly upregulated, but that both 10 nM and 1 nM of 17α-E2 co-treatment suppressed SCD1 protein levels to nearly half the level observed in the vehicle-treated controls (Fig. 6I,J). The effects of 17α-E2 on SCD1 expression in activated HSCs were unanticipated and provides additional evidence that 17α-E2 almost certainly curtails mechanisms of liver disease through direct actions in HSCs, which is suggestive of ERβ-dependency^47,48^.

**Figure 6.**
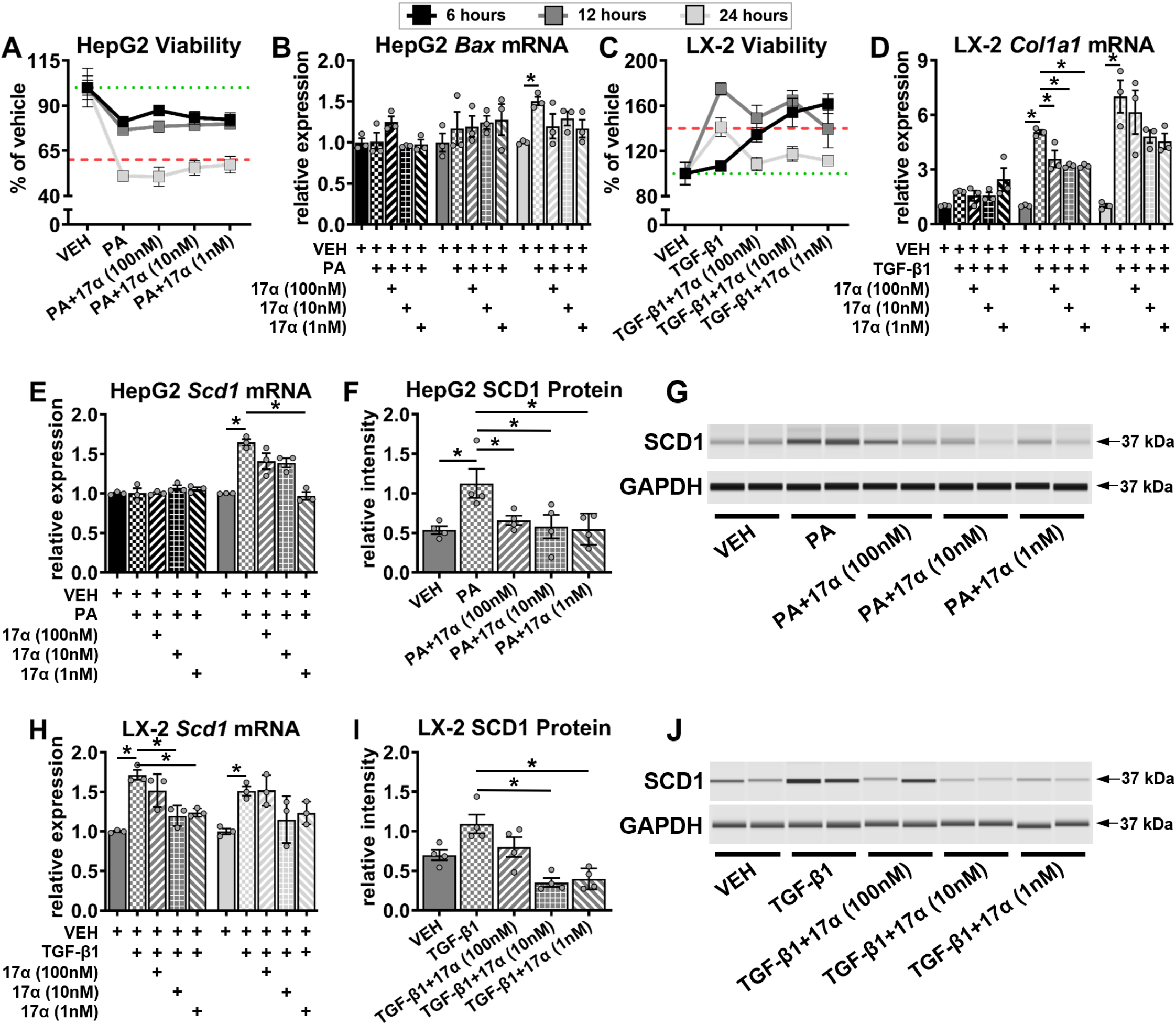
17α-E2 suppresses SCD1 expression in hepatocytes and HSCs *in vitro*. (A) HepG2 cell viability [n=3/treatment/timepoint] and (B) HepG2 *Bax* mRNA [n=3/treatment/timepoint] following 6, 12, and 24 hours of treatment with VEH, VEH+PA (0.5 mM), or VEH+PA+17α-E2 (100 nM, 10 nM, 1 nM). (C) LX-2 cell viability [n=3/treatment/timepoint] and (D) LX-2 *Colla1* mRNA [n=3/treatment/timepoint] following 6, 12, and 24 hours of treatment with VEH, VEH+TGF-β1 (5 ng/ml), or VEH+TGF-β1+17α-E2. (E) HepG2 *Scd1* mRNA [n=3/treatment/timepoint] following 6 and 12 hours of treatment with VEH, VEH+PA, or VEH+PA+17α-E2. (F) HepG2 SCD1 protein [n=4/treatment] following 12 hours of treatment with VEH, VEH+PA, or VEH+PA+17α-E2. (G) Representative immunoblots of SCD1 and GAPDH in HepG2 cells following 12 hours of treatment with VEH, VEH+PA, or VEH+PA+17α-E2. (H) LX-2 *Scd1* mRNA [n=3/treatment/timepoint] following 12 and 24 hours of treatment with VEH, VEH+TGF-β1, or VEH+TGF-β1+17α-E2. (I) LX-2 SCD1 protein [n=4/treatment] following 12 hours of treatment with VEH, VEH+TGF-β1, or VEH+TGF-β1+17α-E2. (J) Representative immunoblots of SCD1 and GAPDH in LX2 cells following 12 hours of treatment with VEH, VEH+TGF-β1, or VEH+TGF-β1+17α-E2. Green dotted lines in panels A and C represent 100% viability, where as the red dash lines represent a 40% loss of viability in panel A and a 40% gain in viability in panel C. All data are presented as mean ± SEM and were analyzed within timepoint by one-way ANOVA (C, D, E, F, H, I) with Tukey post-hoc comparisons. Statistics were not performed on data shown in panels A & C. *p < 0.05. We did not indicate statistical differences between VEH and 17α-E2 treatment groups for purposes of visual clarity.

## DISCUSSION

Previous work has established that 17α-E2 administration ameliorates metabolic dysfunction in obese and aged male mice^25–29^, which we surmise underlies its lifespan-extending effects^20–22^. Female mice are generally unresponsive to 17α-E2 treatment^30–34,36,37^ unless ovariectomized^38^, which suggests 17α-E2 could elicit benefits in female mice in the context of established metabolic dysfunction; although this has not previously been evaluated. We recently reported that the majority of health benefits attributed to 17α-E2 treatment are ERα-dependent^27^. However, we have also shown that 17α-E2 attenuates fibrogenesis in liver^46^, which raises the possibility that 17α-E2 signals directly in ERβ-expressing HSCs^47,48^. Therefore, the studies outlined in this report sought to determine if ERβ plays a role in 17α-E2-mediated benefits on systemic metabolic parameters in the context of obesity in both sexes. We also evaluated how 17α-E2 treatment modulates markers of hepatic steatosis and fibrosis and their interactions with ERβ ablation. Lastly, we also assessed how 17α-E2 treatment alters hepatocyte and HSC responsiveness to metabolic challenges *in vitro*. Several anticipated, and a few unanticipated, outcomes were observed through these studies.

When 17α-E2 treatment was initiated our first objective was to determine if the global ablation of ERβ attenuated 17α-E2-mediated benefits on systemic metabolism in a sex-specific manner. As expected, the ablation of ERβ had little effect on the ability of 17α-E2 to reduce body mass or adiposity in male mice. 17α-E2 was also equally effective at improving hyperinsulinemia and insulin sensitivity in male WT and ERβKO mice in our study. Notably, insulin sensitivity was essentially identical between the male WT LFD, WT HFD+17α-E2, and ERβKO HFD+17α-E2 groups, despite the fact that the HFD-fed groups weighed nearly 10 grams more than the mice receiving LFD. This indicates that 17α-E2 restores metabolic flexibility in the presence of obesity in male mice, which supports our previous findings^27^. Although glucose response curves during the ITT were similar between HFD-fed controls and 17α-E2 treated animals, fasting glucose levels at baseline were significantly lower in animals receiving 17α-E2, which further supports the notion that 17α-E2 treated animals are metabolically healthier than HFD-fed controls.

Female responsiveness to 17α-E2 was one of the most unexpected and important findings from these studies. As addressed previously, female mice are generally unresponsive to 17α-E2 treatment^30–34,36,37^, which we speculate is related to the inherent metabolic advantage provided by endogenous 17β-E2^27,58^. However, 17α-E2 treatment reduced mass and adiposity in female WT mice in the current study, which is likely related to chronic HFD exposure that induced sufficient obesity for 17α-E2 to render health benefits. Interestingly, 17α-E2-mediated effects on body mass and adiposity were attenuated in female ERβKO mice, thereby suggesting that ERβ at least partially controls 17α-E2 responsiveness in obese female mice. Additionally, 17α-E2 treatment reduced calorie consumption in female WT mice, which was a novel finding, but this effect was attenuated in female ERβKO mice. 17α-E2 treatment also only improved hyperinsulinemia and insulin sensitivity in female WT, and not ERβKO, mice. Collectively, these observations certainly suggest that ERβ is required for 17α-E2 to improve metabolic parameters in obese female mice. However, it should be noted that despite being obese, female mice in our study were still fairly resilient to the metabolic detriments of chronic HFD exposure as compared to their male counterparts. For example, the initial glucose levels seen during the ITTs were significantly higher in males than females, and fasting insulin levels were nearly three-fold greater in males than females; both of which indicate that the female mice in our study were metabolically flexible. It is also important to mention that prior reports indicate that the ablation of ERβ can actually improve metabolic outcomes in female mice^59,60^, so it is possible that female ERβKO mice are unresponsive to 17α-E2 treatment because they were metabolically healthier than the female WT mice. Future studies will be needed to unravel this possibility.

Since 17α-E2 is known to suppress mechanisms that promote chronic liver disease^25,27,46^, we next evaluated whether ERβ is required for any of these benefits to occur. In alignment with our prior studies, 17α-E2 prevented and/or reversed hepatomegaly and hepatic lipid accumulation in both male WT and ERβKO mice. Interestingly, improvements in steatosis were not accompanied by changes in liver macrophage content and polarity in either male genotype. We surmise that the lack of changes in macrophage outcomes stems from the fact that liver injury was still fairly minimal in this study, as evidenced by the magnitude of liver fibrosis. We have previously reported that 17α-E2 treatment reduced hepatic total macrophage and M1 proinflammatory macrophage content, while concomitantly increasing M2 anti-inflammatory macrophages in a more robust model of liver injury^46^; therefore, we speculate that improvement would become apparent in models with sufficient injury. This notion is supported by our finding that 17α-E2 treatment significantly suppressed liver *TNFα* transcripts in both male WT and ERβKO mice, which is known to play an important role in sustaining the pro-inflammatory cytokine loop during liver inflammation and the progression toward advanced liver disease^53^.

17α-E2 treatment prevented and/or reversed hepatic lipid accumulation in female mice independently of ERβ in the current study. This is an interesting observation because it suggests that 17α-E2-mediated effects on body mass and adiposity occur through different mechanisms than those that control hepatic lipid metabolism in female mice. In contrast to male mice, both female genotypes were found to have reductions in liver inflammation and pro-inflammatory macrophage content with 17α-E2 treatment. However, it should be noted that liver inflammation was minimal in this study and the observed changes were subtle. We surmise that this stems from the fact that female mice remained relatively healthy throughout the interventional period despite being obese. As alluded to previously, additional studies in models of advanced liver disease (*e.g*. NASH) will provide greater insight into the efficacy of 17α-E2 for the treatment of severe liver pathology in female mice.

Given our previous report showing that 17α-E2 treatment can suppress the production of hepatic SCD1^46^, the rate-limiting enzyme for monounsaturated fatty acid formation that is closely associated with liver steatosis^61–63^ and fibrosis^51,64^, we also evaluated it in the current study. As expected, 17α-E2 treatment robustly suppressed hepatic SCD1 in male WT mice, but this effect was attenuated by the ablation of ERβ. We also found that TGF-β1, the master regulator of liver fibrosis^49,50^, was suppressed by 17α-E2 treatment in male WT, but not ERβKO, mice. We perceive these to be important findings because they suggest that 17α-E2 likely elicits benefits through ERβ-expressing cells within the liver, which is a unique feature of HSCs^47,48^. Interestingly, female mice receiving 17α-E2 responded similarly to male mice with regard to hepatic SCD1 expression, with WT mice responding more robustly, although TGF-β1 production was unchanged in female mice of either genotype. These findings indicate that 17α-E2 can also suppress SCD1 in female mice through actions in hepatocytes, but that 17α-E2 is either incapable of modulating pro-fibrotic mechanisms in female liver, or the female mice in these studies were too metabolically healthy to develop sufficient liver pathology for 17α-E2 to elicit benefits. We speculate the latter is more likely given the magnitude of disease burden displayed by female mice in our studies.

In an effort to confirm our suspicion that 17α-E2 elicits benefits in both hepatocytes and HSCs, we then evaluated the effects of 17α-E2 in cultured HepG2 and LX-2 cells had been challenged with PA and TGF-β1, respectively. After establishing appropriate culture conditions we quickly determined that 17α-E2 indeed suppressed SCD1 at the mRNA and protein levels in both hepatocytes and HSCs. We anticipated that 17α-E2 would attenuate SCD1 in stressed hepatocytes due to our previous reports^25,27,46^, but the suppression in HSCs was unexpected. These observations provide clear evidence that 17α-E2 almost certainly curtails mechanisms of liver disease through direct actions not only in hepatocytes, but also HSCs, the latter of which is suggestive of ERβ-dependency^47,48^. The ability of 17α-E2 to elicit benefits in both cell-types, independently, is very important because it is now recognized that novel therapies for treating NASH and/or liver fibrosis must target multiple pathways through several cell-types for successful translation into clinical trials^65–67^.

There are a few notable caveats to the current studies that should be acknowledged. First, a minor limitation is that the female mice we studied were still relatively healthy, despite being. This prevented us from making definitive conclusions regarding the role that ERβ plays in regulating 17α-E2-mediated effects on systemic metabolism and hepatic profibrotic mechanisms in female mice. Since HFD feeding does not recapitulate human liver disease^55^, future studies utilizing a combination approach of western diet and CCl_4_ administration should be undertaken to determine if 17α-E2 also elicits benefits in a disease state that more closely resembles human NASH and liver fibrosis^68^. We speculate that 17α-E2 will elicit even greater benefits in models of human NASH and liver fibrosis. Another minor limitation of the animal studies was that the global ablation of ERβ has been reported to improve metabolic parameters in female mice^59,60^, which partially limited our ability to determine how ERβ is involved in metabolic regulation by 17α-E2 due to the mice being resilient to the metabolic challenge. Future studies utilizing hepatocyte- and HSC-specific deletions of ERα or ERβ will provide tremendous insight into the effects of 17α-E2 on chronic liver disease.

In summary, the data presented herein are the first to show that ERβ is not required for 17α-E2 to improve systemic metabolic parameters in male mice. We also show that metabolically challenged female mice are responsive to 17α-E2 treatment and that ERβ appears to at least partially mediate these effects. 17α-E2 was again found to improve a variety of parameters related to liver steatosis and fibrosis, particularly in male mice. Our most important discovery was that 17α-E2 directly elicits benefits through direct actions in hepatocytes and HSCs, which is an important characteristic for novel therapies aimed at treating chronic liver diseases. Our current studies provide critical insight into the how 17α-E2 may have therapeutic potential for the treatment of chronic liver diseases.

## METHODS

### Animal diets

Control animals receiving standard chow diet (LFD) were provided TestDiet 58YP (66.4% CHO, 20.5% PRO, 13.1% FAT). Animals receiving HFD were provided TestDiet 58V8 (35.5% CHO, 18.3% PRO, 45.7% FAT) and animal receiving HFD with 17α-E2 (HFD+17α) were provided TestDiet 58V8 supplemented with 14.4 ppm of 17α-E2 (Steraloids, Newport, RI, USA) during the manufacturing process. All diets are identical to those used in prior studies with ERα knockout (ERαKO) mice^27^.

### Animal experiments

Heterozygous ERβ knockout (ERβKO) mice were acquired from the laboratory of Dr. Jan-Åke Gustafsson (Karolinska Institute, Stockholm, SE). Experimental mice were generated by breeding heterozygous ERβKO mice within the Oklahoma City VA Health Care System vivarium so that wild-type (WT) and homozygous ERβKO littermates of both sexes could be enrolled in our study. At weaning, experimental mice were group-housed by sex and fed standard chow (LFD) until 12 weeks of age. At 12 weeks of age all mice, excluding the WT LFD controls, were fed HFD for 39 weeks (9 months) to induce obesity and metabolic perturbations prior to study initiation. The age-matched WT LFD control mice were evaluated in parallel as a healthy-weight reference group. Two weeks prior to study initiation, all mice were individually housed with ISO cotton pad bedding, cardboard enrichment tubes, and nestlets at 22 ± 0.5°C on a 12:12-hour light-dark cycle. Unless otherwise noted, all mice had ad libitum access to food and water throughout the experimental timeframe. At the conclusion of the 39-week fattening period, all mice receiving HFD were randomized within genotype by baseline body mass, fat mass, calorie intake, and fasting insulin into HFD or HFD+17α treatment groups for a 10-week intervention. Body mass and calorie intake were assessed daily for the first 4 weeks, followed by body mass and body composition (EchoMRI, Houston, TX, USA) on a weekly basis. During the ninth week of treatment, mice were fasted and glucose tolerance was assessed. During the tenth week of treatment, mice were fasted and insulin tolerance was assessed. At the conclusion of the interventional period, mice were anesthetized with isoflurane following a 5-6 hour fast and euthanized by exsanguination via cardiac puncture. Blood was collected into EDTA-lined tubes, plasma was collected and frozen, and the mice were then perfused with ice-cold 1X PBS prior to tissues being excised, weighed, flash-frozen and store at −80°C unless otherwise noted. Following excision, small pieces of liver in the portal triad region were dissected and fixed in 4% PFA in preparation for paraffin- or cryo-embedding for future analyses. All animal procedures were reviewed and approved by the Institutional Animal Care and Use Committee of the Oklahoma City VA Health Care System.

### *In vivo* metabolic analyses

All experiments requiring fasting conditions were performed in the afternoon, 5-6 hours following the removal of food at the beginning of the light-cycle for reasons outlined elsewhere^69^. To ensure fasting conditions, mice were transferred to clean cages containing ISO cotton padding and clean cardboard enrichment tubes. Non-terminal blood was collected via tail snip. Glucose tolerance tests (GTT) were performed following the administration of a filtered dextrose (1 g/kg body mass) solution via IP injection^70^. Insulin tolerance tests (ITT) were performed following the administration of a filtered insulin (0.75 mU/g body mass; Novolin-R, Novo Nordisk, Bagsvaerd, DK) solution via IP injection^71^. Blood glucose was measured immediately pre-injection (time 0) and at 15, 30, 45, 60, 90, and 120 minutes post-injection during the GTT and ITT. The area under curve (AUC) for each animal during both the GTT and ITT were also calculated and presented as the average for each group. Blood glucose levels were determined using Accu-Chek Aviva Plus glucometers (Roche, Basel, CH). Fasting insulin levels from blood collected at baseline and during the terminal harvest were evaluated using a Mouse Ultrasensitive Insulin ELISA from Alpco (Salem, NH, USA).

### Liver triglyceride analyses

Liver samples (~100 mg) were homogenized on ice for 60 seconds in 10X (v/w) RIPA Buffer (Cell Signaling, Danvers, MA, USA) with phosphatase and protease inhibitors (Boston BioProducts, Boston, MA, USA). Total lipid was extracted from 100 ul of homogenate using the Folch method^72^. Samples were dried under nitrogen gas at room temperature prior to being resuspended in 100μl of tert-butyl alcohol-methanol-Triton X-100 solution (3:1:1). Triglycerides (TG) were quantified spectrophotometrically using Free Glycerol & Triglyceride agents (Sigma-Aldrich, St. Louis, MO, USA) as previously described^73^. The remaining liver homogenate was used for western blotting and TGF-β1 quantification as outlined below.

### Liver histology and pathology assessments

Liver samples were fixed in 4% PFA for 24 hours, transferred to 1X PBS for 48 hours, and then transferred to 70% EtOH until paraffin embedding occurred. H&E and Masson’s trichrome staining were performed by the OMRF Imaging and Histology Core Facility using established protocols. Images of H&E and trichrome stained slides were taken on an Olympus CX43 microscope and were evaluated by two clinical pathologists who were blinded to the treatment groups as previously described^46^. NAFLD activity scores (NAS) and fibrosis staging were determined according to NASH Clinical Research Network standards^74,75^.

### Liver immunofluorescence analyses

Cryo-embedded liver samples were sectioned (10 μm) and stained with primary antibodies against EGF-like module-containing mucin-like hormone receptor-like 1 (F4/80; total macrophages; USBiological Life Sciences, Salem, MA USA; 1:250), integrin, alpha X (CD11c; M1, pro-inflammatory macrophages; Invitrogen; 1:300), and mannose receptor (CD206; M2, anti-inflammatory macrophages; Cell Signaling; 1:500) as previously described^46,76^. Secondary antibodies used included goat anti-Armenian hamster IgG Alexa Fluor 594 (Jackson ImmunoResearch Laboratories, West Grove, PA, USA; 1:500), goat anti-chicken IgG Alexa Fluor 647 (Jackson ImmunoResearch Laboratories; 1:500), and goat anti-rabbit IgG Alexa Fluor 488 (Jackson ImmunoResearch Laboratories; 1:500). Sections were mounted in Prolong Diamond Mounting Medium with DAPI (Abcam) and images were acquired using a Leica 3D Thunder scope from three non-intersecting fields per mouse. Intensity of fluorescence was measured as percent of total area using Image J after each image had its background intensity subtracted out.

### Quantitative real-time PCR

Total RNA was extracted using Trizol (Life Technologies, Carlsbad, CA, USA) and RNA (2μg) was reverse transcribed to cDNA with the High-Capacity cDNA Reverse Transcription kit (Applied Biosystems, Foster City, CA, USA). Real-time PCR was performed in a QuantStudio 12K Flex Real Time PCR System (ThermoFisher Scientific) using TaqMan™ Gene Expression Master Mix (ThermoFisher Scientific) and predesigned gene expression assays with FAM probes from Integrated DNA Technologies (Skokie, Illinois, USA). Target gene expression for mouse (liver tissue) sterol regulatory element-binding protein 1 (*Srebp1*), fatty acid synthase (*Fasn*), acetyl-CoA carboxylase 1 (*Acc1*), and tumor necrosis factor α (*TNFα*) are expressed as 2^-ΔΔCT^ by the comparative CT method^77^ and normalized to the expression of TATA-box binding protein (*Tbp*). Target gene expression for human (cultured cells) bcl-2-like protein 4 (*Bax*), collagen, type 1, alpha 1 (*Col1a1*), and *Scd1* are expressed as 2^-ΔΔCT^ by the comparative CT method and normalized to the expression of glyceraldehyde-3-phosphate dehydrogenase (*Gapdh*).

### Western blot analyses

Liver homogenates not used for the liver triglyceride analyses were spun at 17,000 rpm for 15 minutes at 4°C and the supernatant was collected. Total protein was quantified using a Pierce BCA kit (ThermoFisher Scientific, Waltham, MA, USA). Proteins were separated on Any kD Criterion TGX Stain-Free gels (Bio-Rad, Hercules, CA, USA) at 75V for 150 minutes in running buffer (Cell Signaling). Protein was then transferred to 0.2 μm pore-size nitrocellulose membranes (Bio-Rad) at 75V for 90 minutes on ice. Both primary antibodies utilized have been commercially validated and include SCD1 (Cell Signaling; 1:1000) and GAPDH (Abcam, Waltham, MA, USA; 1:2500). Primary antibody detection was performed with IRDye 800CW Infrared Rabbit antibody (LI-COR Biotechnology, Lincoln, NE, USA) at a concentration of 1:15000. Imaging was done on an Odyssey Fc Imaging System (LI-COR Biotechnology) protein quantification was performed using Image Studio Software (LI-COR Biotechnology).

### Liver TGF-β1 quantification

Twenty microliters of supernatant from liver homogenates were diluted with TGF-β1 ELISA (Abcam) assay buffer (180 ul), which was then digested with 1N HCl (20 ul) for 60 minutes at room temperature. Samples were then neutralized with 1N NaOH (20 ul) and were further diluted with ELISA assay buffer to a total volume of 1200 ul (1:60 dilution). The samples were then evaluated according manufacturer instructions as we described previously^46^. TGF-β1 concentrations were normalized to total protein and expressed as ng/mg protein.

### *In vitro* experiments

Immortalized human HepG2 cells (ATCC, Manassas, VA, USA) were cultured in phenol-free DMEM (Sigma-Aldrich) containing 10% FBS (Gibco, Grand Island, NY, USA) and 1% penicillin-streptomycin (P/S; Gibco) at 37° C with 5% CO2. When cells reached 80-90% confluency they were washed with PBS and cultured in serum-free DMEM with 1% P/S overnight. Cells were then treated with vehicle (VEH: DMSO; Sigma-Aldrich) or 17α-E2 (100nM, 10nM, 1nM; Steraloids) in VEH for 60 minutes prior to the addition of palmitic acid (PA; 0.5mM; Cayman Chemical, Ann Arbor, MI, USA). At 6, 12, & 24 hours post-treatment, cells were washed with PBS and evaluated for cell viability using an MTT Assay Kit (Abcam), or were harvested for mRNA and protein assessment. Immortalized human LX-2 cells (MilliporeSigma, Burlington, VT, USA) were cultured in phenol-free DMEM (Sigma-Aldrich) containing 2% FBS (Gibco, Grand Island, NY, USA), 1% penicillin-streptomycin (P/S; Gibco), and 4 mM L-glutamine (L-Glu, Gibco) at 37° C with 5% CO2. When cells reached 80-90% confluency they were washed with PBS and cultured in serum-free DMEM with 0.1% BSA overnight. Cells were then treated with VEH or 17α-E2 (100nM, 10nM, 1nM) in VEH for 60 minutes prior to the addition of recombinant human TGF-β1 (5 ng/ml; R&D Systems, Minneapolis, MN, USA). At 6, 12, & 24 hours post-treatment, cells were washed with PBS and evaluated for cell viability as describe above, or were harvested for mRNA and protein assessment. Total RNA from cultured cells was extracted and processed identically to that of mouse liver described above. Total protein from cultured cells was extracted and processed identically to that of mouse liver described above, with the exception that western blots were ran on the Jess SimpleWestern System (R&D Systems) as described previously^78^ due to low protein abundance. Both primary antibodies utilized have been commercially validated and include SCD1 (Abcam; 1:25) and GAPDH (Abcam; 1:50).

### Statistical analyses

Results are presented as mean ± SEM with *p* values less than 0.05 considered significantly different. Analyses of differences between groups were performed by two-way ANOVA, repeated measures two-way ANOVA, or one-way ANOVA with Tukey post-hoc comparisons where appropriate using GraphPad Prism Software, Version 9.

## ACKNOWLEDGEMENTS

We thank Drs. Per Antonson and Jan-Åke Gustafsson from the Karolinska Institute for generously providing heterozygous ERβKO mice which enabled us to generate a colony for these studies. This work was supported by the National Institutes of Health (T32 AG052363 to S.N.M. & M.P.B., R00 AG056662 to S.L., and R00 AG051661 & R01 AG070035 to M.B.S.) and the US Department of Veterans Affairs (I01 BX005592 to B.F.M. and Pilot Research Funding to M.B.S.).

## AUTHOR CONTRIBUTIONS

S.A.M., S.N.M., and M.B.S. conceived the project and designed the experiments. S.A.M. and S.N.M. performed the experiments with contributions from C.v.d.L., R.S., M.K., W.L., M.P.B., S.L., and B.F.M. S.A.M., S.N.M., and M.B.S. created figures and performed statistical analyses. S.A.M., C.v.d.L., and M.B.S. wrote the manuscript and all authors edited and approved the final manuscript.

## DATA AVAILABILITY STATEMENT

The datasets generated through this work are available upon reasonable request from the corresponding author.

## COMPLETING INTERESTS

The authors declare no conflicts or competing interests.

## FIGURE LEGENDS

**Supplemental Figure 1.**
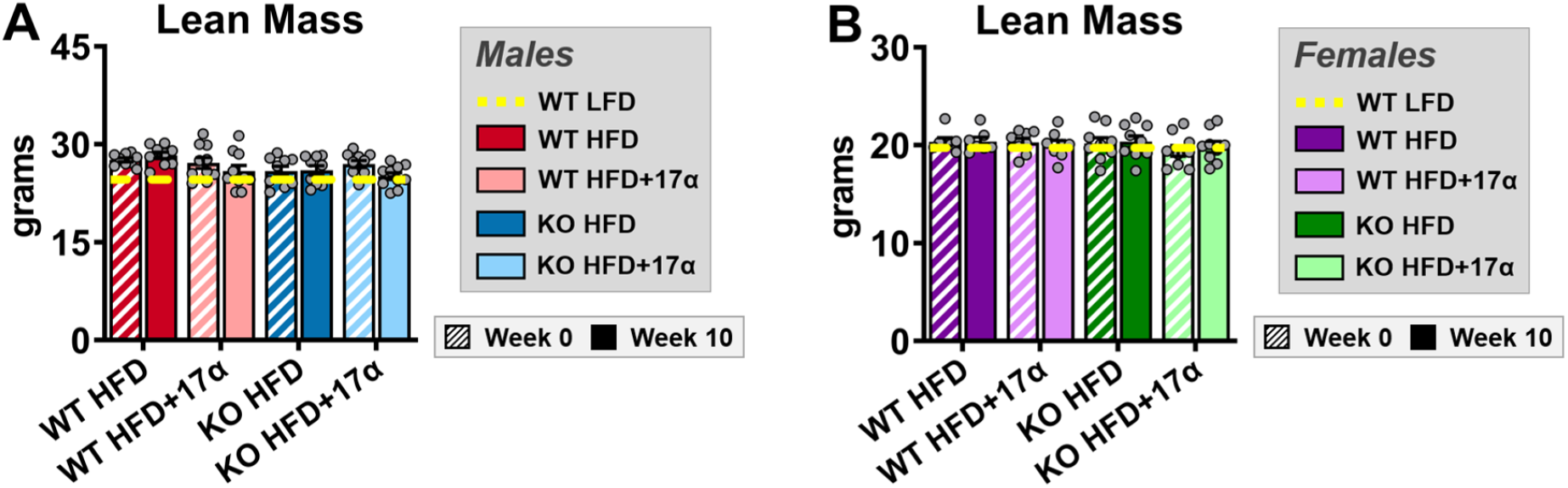
Neither ERβ ablation, nor 17α-E2 treatment, affects lean mass in obese mice of either sex. (A) Lean mass at baseline (week 0; striped) and week 10 (solid) in male WT and ERβKO mice [n=9-10/group/timepoint]. (B) Lean mass at baseline (week 0; striped) and week 10 (solid) in female WT and ERβKO mice [n=7-9/group/timepoint]. Age-matched, WT, LFD-fed mice were also evaluated as a normal-weight reference group and their corresponding means for both sexes are depicted as dashed yellow lines [n=9/group/timepoint]. All data are presented as mean ± SEM and were analyzed within sex by two-way repeated measures ANOVA with Tukey post-hoc comparisons.

**Supplemental Figure 2.**
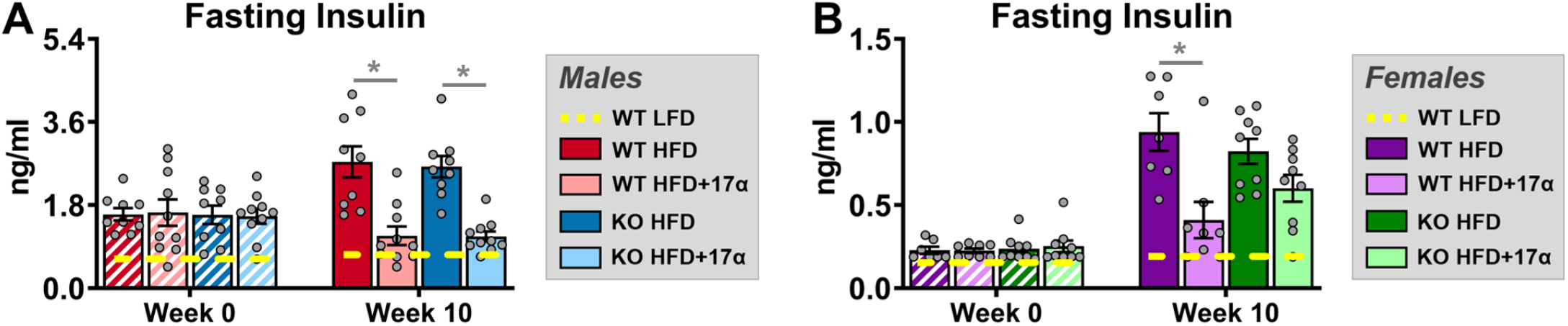
ERβ partially mediates 17α-E2 effects on fasting insulin in obese female, but not male, mice. (A) Fasting insulin at baseline (week 0; striped) and week 10 (solid) in male WT and ERβKO mice [n=9-10/group/timepoint]. (B) Fasting insulin at baseline (week 0; striped) and week 10 (solid) in female WT and ERβKO mice [n=7-9/group/timepoint]. Age-matched, WT, LFD-fed mice were also evaluated as a normal-weight reference group and their corresponding means for both sexes are depicted as dashed yellow lines [n=9/group/timepoint]. All data are presented as mean ± SEM and were analyzed within sex by two-way repeated measures ANOVA with Tukey post-hoc comparisons. * represents differences within genotypes across treatment groups at each timepoint. *p < 0.05. We did not indicate statistical differences between week 0 and week 10 for purposes of visual clarity.

**Supplemental Figure 3.**
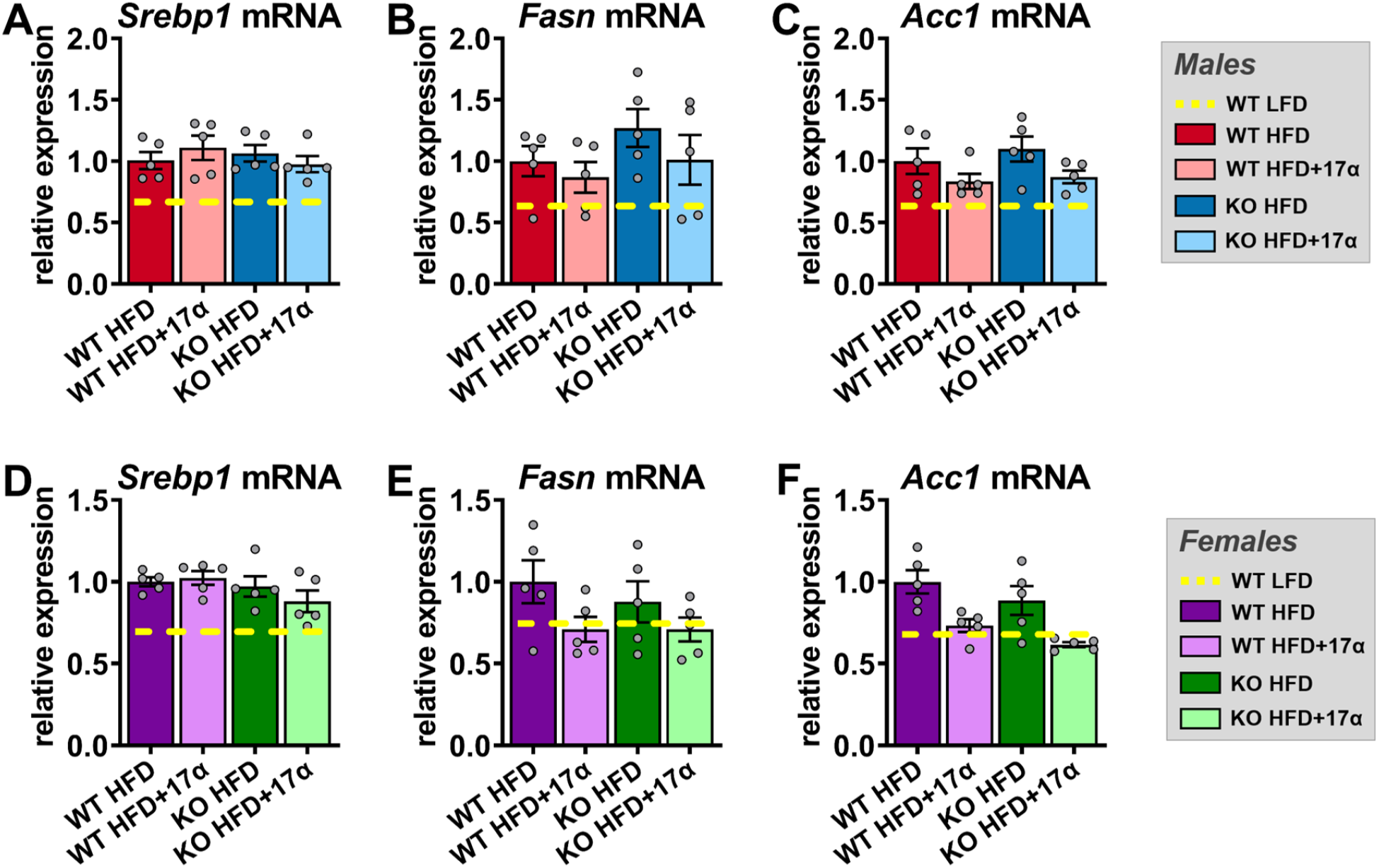
Neither ERβ ablation, nor 17α-E2 treatment, effects hepatic transcriptional markers of lipogenesis in obese mice of either sex. (A) *Srebp1* mRNA [n=5/group], (B) *Fasn* mRNA [n=5/group], and (C) *Acc1* mRNA [n=5/group] in liver from male WT and ERβKO mice. (D) *Srebp1* mRNA [n=5/group], (E) *Fasn* mRNA [n=5/group], and (F) *Acc1* mRNA [n=5/group] in liver from female WT and ERβKO mice. Age-matched, WT, LFD-fed mice were also evaluated as a normal-weight reference group and their corresponding means for both sexes are depicted as dashed yellow lines [n=5/group]. All data are presented as mean ± SEM and were analyzed within sex by two-way ANOVA with Tukey post-hoc comparisons.

**Supplemental Figure 4.**
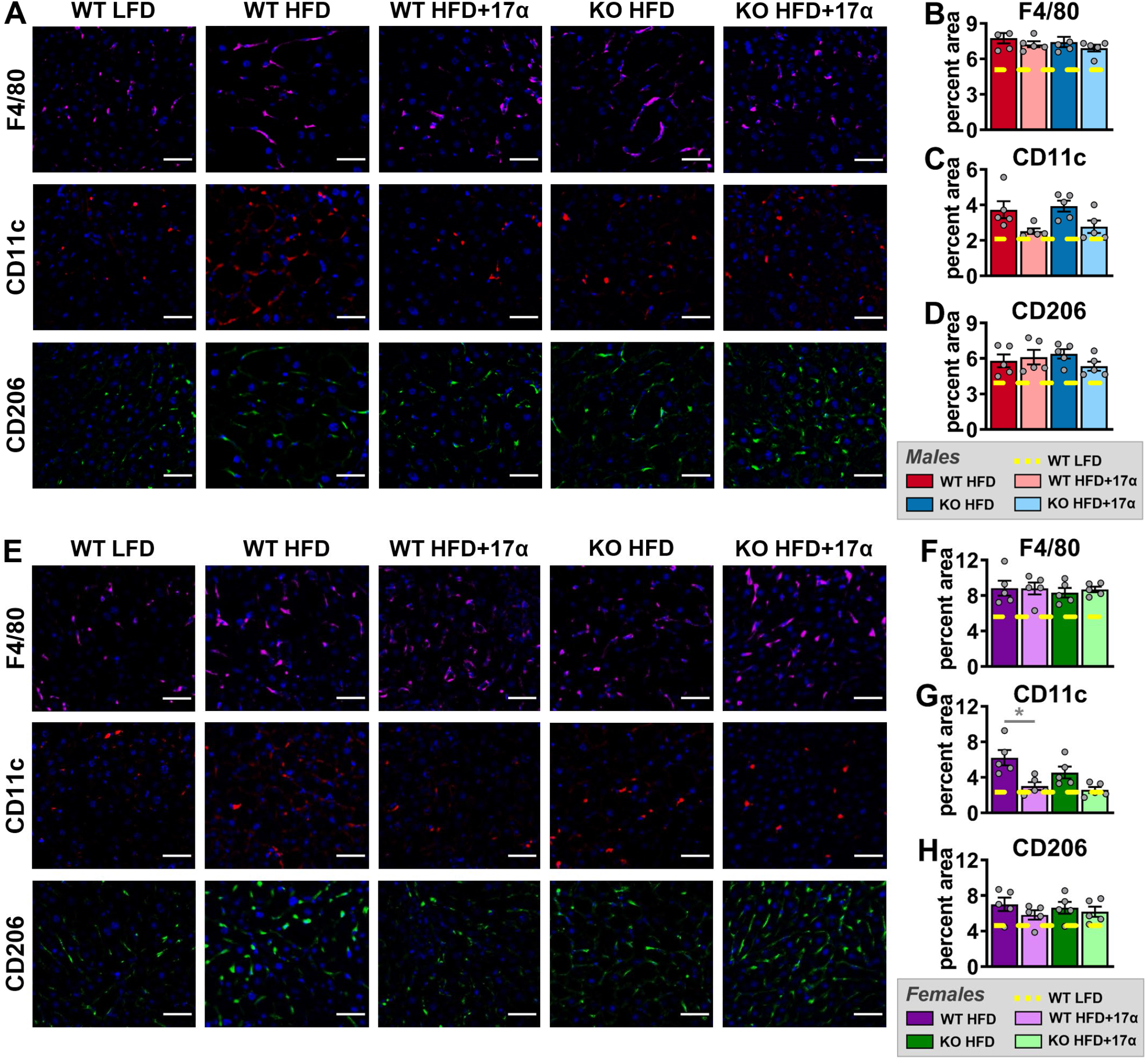
17α-E2 attenuates proinflammatory macrophage responses in female, but not male, mice in a ERβ-dependent manner. (A) Representative immunofluorescence images of F4/80 (total macrophages), CD11c (M1, pro-inflammatory macrophages), and CD206 (M2, anti-inflammatory macrophages) in liver from male WT and ERβKO mice (magnification = 320X; scale bar = 50 μm). (B) Percent area for F4/80 [n=5/group], (C) Percent area for CD11c [n=5/group], and (D) Percent area for CD206 [n=5/group] in liver from male WT and ERβKO mice. (E) Representative immunofluorescence images of F4/80 (total macrophages), CD11c (M1, pro-inflammatory macrophages), and CD206 (M2, anti-inflammatory macrophages) in liver from female WT and ERβKO mice (magnification = 320X; scale bar = 50 μm). (F) Percent area for F4/80 [n=5/group], (G) Percent area for CD11c [n=5/group], and (H) Percent area for CD206 [n=5/group] in liver from female WT and ERβKO mice. Age-matched, WT, LFD-fed mice were also evaluated as a normal-weight reference group and their corresponding means for both sexes are depicted as dashed yellow lines [n=5/group]. All data are presented as mean ± SEM and were analyzed within sex by two-way ANOVA with Tukey post-hoc comparisons. * represents differences within genotypes across treatment groups. *p < 0.05.

**Supplemental Figure 5.**
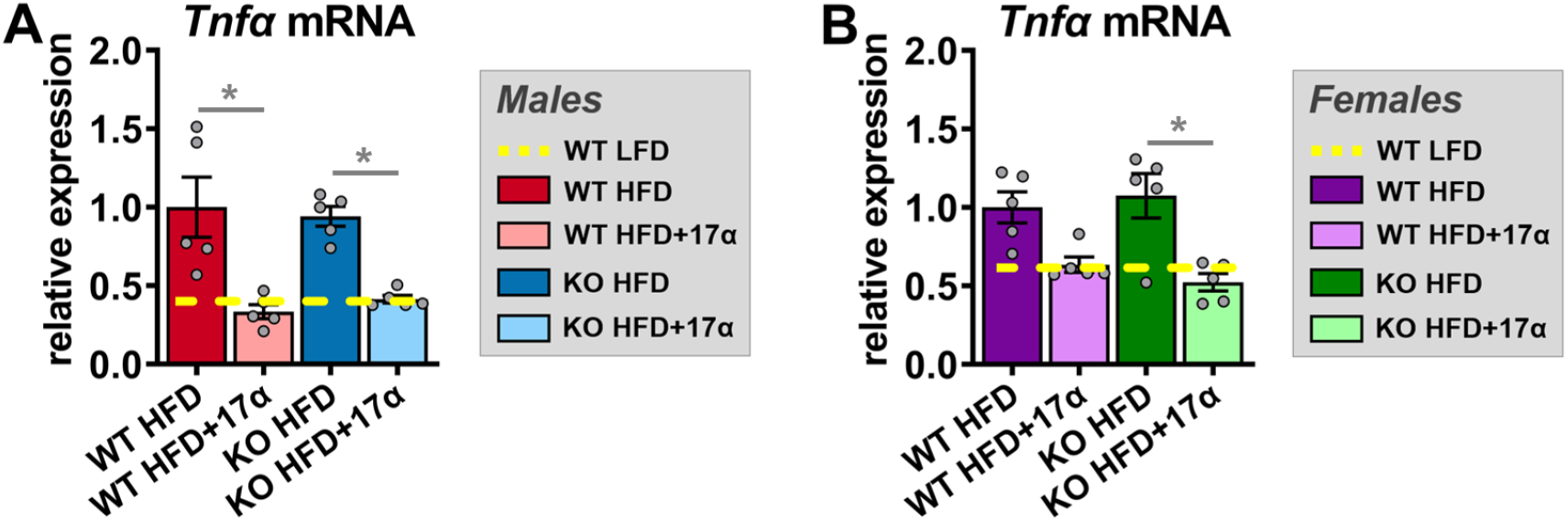
17α-E2 suppresses hepatic tumor necrosis factor α transcripts in both sexes in an ERβ-independent manner. (A) *TNFα* mRNA in liver from male WT and ERβKO mice [n=5/group]. (B) *TNFα* mRNA in liver from female WT and ERβKO mice [n=5/group]. Age-matched, WT, LFD-fed mice were also evaluated as a normal-weight reference group and their corresponding means for both sexes are depicted as dashed yellow lines [n=5/group]. All data are presented as mean ± SEM and were analyzed within sex by two-way ANOVA with Tukey post-hoc comparisons. * represents differences within genotypes across treatment groups. *p < 0.05.

